# A Canal-Associated Neuron cAMP signalling pathway that regulates *C. elegans* larval development

**DOI:** 10.1101/733618

**Authors:** Jason Chien, Fred W. Wolf, Sarah Grosche, Nebeyu Yosef, Gian Garriga, Catarina Mörck

## Abstract

*Caenorhabditis elegans* larval development requires the function of the two Canal-Associated Neurons (CANs): killing the CANs by laser microsurgery or disrupting their development by mutating the gene *ceh-10* results in early larval arrest. How these cells promote larval development, however, remains a mystery. In screens for mutations that bypass CAN function, we identified the gene *kin-29,* which encodes a member of the Salt-Inducible Kinase (SIK) family and a component of a conserved pathway that regulates various *C. elegans* phenotypes. Like *kin-29* loss, gain-of-function mutations in genes that may act upstream of *kin-29* or growth in cyclic-AMP analogs bypassed *ceh-10* larval arrest, suggesting that a conserved adenylyl cyclase/PKA pathway inhibits KIN-29 to promote larval development and that loss of CAN function results in dysregulation of KIN-29 and larval arrest. The adenylyl cyclase ACY-2 mediates CAN-dependent larval development: *acy-2* mutant larvae arrested development with a similar phenotype to *ceh-10* mutants, and the arrest phenotype was suppressed by mutations in *kin-29*. ACY-2 is predominantly expressed in the CANs, and we provide evidence that the *acy-2* functions in the CANs to promote larval development. By contrast, cell-specific expression experiments suggest that *kin-29* acts in both the hypodermis and neurons, but not in the CANs. Based on our findings, we propose that cAMP produced by ACY-2 in the CANs acts in neighboring neurons and hypodermal cells where it activates PKA and inhibits KIN-29 to promote larval development. We discuss how this conserved pathway could be partitioned between two cells.

## INTRODUCTION

The nematode *Caenorhabditis elegans* (*C. elegans*) requires only three neurons for survival: the M4 motor neuron and the two Canal-Associated Neurons (CANs). The M4 neuron is located in the pharynx, the *C. elegans* feeding organ, and is required for peristaltic movements that move food along the pharynx (AVERY AND HORVITZ 1987; AVERY AND HORVITZ 1989). The CANs are two bilaterally symmetric neurons that are born in the head and migrate posteriorly to the middle of the worm during embryogenesis. After the CANs have completed their migration, each neuron extends two axons: one axon grows anteriorly towards the head, and the other grows posteriorly towards the tail (WHITE *et al.* 1976; WU *et al.* 2011). If the CANs are killed by laser microsurgery or if the neurons fail to differentiate, the worms arrest their development early in larval development (FORRESTER AND GARRIGA 1997; FORRESTER *et al.* 1998). How the CANs regulate larval development is unknown.

Phenotypic analysis of mutants with CAN defects also reveals their role in larval development. The CANs express two differentiation markers, the homeodomain transcription factors CEH-10 and CEH-23 (WANG *et al.* 1993; SVENDSEN AND MCGHEE 1995). Loss of *ceh-10* also results in larval arrest, which is thought to result from the failure of the CANs to differentiate (FORRESTER AND GARRIGA 1997; FORRESTER *et al.* 1998). The posteriorly directed migrations of many cells and growth cones require the gene *vab-8* (<u>v</u>ariable <u>ab</u>normal) (WIGHTMAN *et al.* 1996; WOLF *et al.* 1998). In *vab-8* null mutants, the CANs fail to migrate posteriorly, and their posterior axons fail to extend or extend a short distance. The posterior body of older *vab-8* mutant larvae and adults becomes thin and develops abnormally. This withered tail (Wit) phenotype is thought to result from the lack of CAN function in the posterior of the mutant animals (WIGHTMAN *et al.* 1996), a hypothesis that is supported by a correlation in different mutants between the severity of the defect in the extension of the CAN posterior axon and the penetrance of the Wit phenotype (FORRESTER AND GARRIGA 1997).

In an attempt to reveal the function of the CANs, we mutagenized *ceh-10* or *vab-8* mutants and screened for mutations that can suppress the mutant larval arrest or Wit phenotypes without suppressing their CAN neuron defects. In our screens, we identified three alleles of *kin-29,* which encodes a serine/threonine kinase that is a member of the Salt-Inducible Kinase (SIK) family involved in the regulation of feeding and fasting states (KOO *et al.* 2005; DENTIN *et al.* 2007; WANG *et al.* 2008; CHOI *et al.* 2011).

The three mammalian SIKs are inhibited by a conserved G-protein Coupled Receptor (GPCR) pathway that activates adenylyl cyclase (ACY) and Protein Kinase A (PKA) (WANG *et al.* 1999; TAKEMORI *et al.* 2002; OKAMOTO *et al.* 2004; VAN DER LINDEN *et al.* 2008). Here we report that mutations that cause an increase of cAMP levels or the activation of PKA rescue the *ceh-10* larval arrest phenotype. We also provide evidence that ACY-2 is the adenylyl cyclase that generates the cAMP necessary for CAN-dependent larval development. ACY-2 is expressed in the CANs and in a few other neurons (KORSWAGEN *et al.* 1998). We found that when expressed in the CANs, *acy-2* partially rescued the mutant *acy-2* larval arrest phenotype. Furthermore, CAN-specific RNAi of *acy-2* induced larval arrest. Together, these findings suggest that ACY-2 produces cAMP in the CANs. To address where KIN-29 acts when CAN function is defective, we performed cell-specific expression experiments that suggest that KIN-29 functions in the hypodermis and neurons, but not in the CANs. Our observations are consistent with a model where cAMP produced by ACY-2 in the CANs negatively regulates KIN-29 in neurons and hypodermal cells to promote larval development. We propose that cAMP diffuses from the CANs through gap junctions to inhibit KIN-29 though PKA to promote proper larval development.

## Materials and Methods

### C. elegans genetics

Worms were cultured as previously described (BRENNER 1974). All strains were maintained at 20°C, unless otherwise noted. The following mutant alleles were used: LGI: *gsa-1(ce94), lin-35(n745), mef-2(gv1)* LGII: *pde-4(ce268)* LGIII: *acy-1(pk1279), ceh-10(gm58), rrf-3(pk1426)* LGIV: *eri-1(mg366)* LGV: *acy-2(pk465), ergo-1(gg100), nre-1(hd20), rde-1(ne219), vab-8(e1017)* LGX: *hda-4(oy59), kin-2(ce179), kin-29(gm112), kin-29(jehm1), kin-29(jehm2), kin-29(gk288), lin-15B(n744), lin-15B(hd126)* Transgenes: *gmIs18[Pceh-23::GFP;pRF4(rol-6su1006)] (ZINOVYEVA AND FORRESTER 2005)*

The double and triple mutants created in the genetic interaction studies were sequenced to confirm that all mutations were present.

### ceh-10 and vab-8 suppressor screens

*ceh-10(gm58)/ht2; gmIs18* or *vab-8(e1017)gmIs18* worms were mutagenized for 4 hours by incubation in 0.05 M ethyl methane sulfonate (EMS). Worms were washed in M9 buffer (22mM KH_2_P0_4,_ 42 mM Na_2_HP0_4_, 85.5 mM NaCl and 1mM MgS0_4_) and placed on a large culture dish. 2 hours later L4 hermaphrodites were transferred to new plates in groups of five worms. Five to six days later F1 progeny were picked individually to new plates and on day 9-13, the F2 progeny were screened for ability to rescue *ceh-10* larval arrest or *vab-8* withered tail phenotype. *gm112* suppressed the *vab-8* Wit phenotype and *jehm1* and *jehm2* suppressed *ceh-10* larval arrest. All suppressors were outcrossed ten times to the wild-type N2 strain.

### Mutant identification

For identification of *kin-29(gm112),* we used a combination of SNP mapping, RNAi interference and sequencing. The Hawaiian isolate CB4856 was used for SNP mapping (WICKS *et al.* 2001), which placed *gm112* between SNPs in the *H01M10.1* and *pccb-1* genes. Genes located between *H01M10.1* and *pccb-1* were tested for suppression of *vab-8* Wit and *ceh-10* larval arrest by feeding worms bacteria expressing double-stranded RNA specific to a single gene. RNAi clones were obtained from the Ahringer RNAi library (KAMATH *et al.* 2003) or the *C. elegans* ORFeome library (RUAL *et al.* 2004) and were verified by sequencing. The experiments were performed as previously described (TIMMONS *et al.* 2001).

RNAi against *kin-29* rescued both *vab-8* Wit and *ceh-10* larval arrest phenotypes. The mutant *kin-29* genes were sequenced by amplifying fragments covering the entire *kin-29* gene by PCR.

### DNA plasmid constructs and transgenic lines

***Pges-1::kin-29cDNA*** was generated by PCR amplifying 3323 bp of the *ges-1* promoter using wild type genomic DNA as template with the following primers: 5’-<u>ctcgag</u>ctaagcttaatgaagtttatttc-3’ (*XhoI* site underlined) and 5’-<u>ggatcc</u>ctgaattcaaagataagatatgt-3’(*BamHI* site underlined). The PCR product was cloned into pCR®2.1-TOPO® (Invitrogen), cut out with *XhoI* and *BamHI* and ligated into pBluescriptKS-. 2468 bp *kin-29*cDNA was amplified from *Pkin-29::kin-29cDNA::GFP* (a kind gift from Piali Sengupta) with the following primers: 5’-<u>ggatcc</u>atggctgcgccacggcggc-3’(*BamHI* site underlined) and 5’-<u>gcggccgct</u>cactccgagctccagcttg-3’(NotI site underlined). The PCR product was cloned into pCR®2.1-TOPO® (Invitrogen), cut out with *BamHI* and *NotI* and ligated into *Pges-1;pBluescriptKS-.* 744 bp of *unc-54* 3’UTR was generated by PCR amplification using wild type genomic DNA as template with the following primers: 5’-<u>gcggccgc</u>catctcgcgcccgtgcctc-3’(*NotI* site underlined) and 5’-<u>gcggccgc</u>aaacagttatgtttggtat-3’ (*NotI* site underlined). The PCR product was cloned into pCR®2.1-TOPO® (Invitrogen), cut out with *NotI* and ligated into *Pges-1::kin-29cDNA;pBluescriptKS-* creating *Pges-1::kin-29cDNA::unc-54 3’UTR.* The plasmid was injected into *ceh-10;kin-29* at 25 ng/μl together with 2 ng/μl *Pmyo-2::mCherry*.

***Pges-1::GFP*** was generated by PCR amplifying 3323 bp of the *ges-1* promoter using wild type genomic DNA as template with the following primers: 5’-<u>ggatcc</u>ctaagcttaatgaagtttatttc-3’ (*BamHI* site underlined) and 5’-<u>ccatgg</u>ctgaattcaaagataagatatgt-3’(*NcoI* site underlined). The PCR product was cloned into pCR®2.1-TOPO® (Invitrogen), cut out with *BamHI* and *NcoI* and ligated into pPD95.77. The plasmid was injected into wild-type worms at 25 ng/μl together with 40 ng/μl *pRF4 (rol-6 su1006)*.

***Phlh-1:: kin-29cDNA*** was generated by PCR amplifying 3052 bp of the *hlh-1* promoter using wild type genomic DNA as template with the following primers: 5’-<u>ctgcag</u>cagaattctgtgaaataagc-3’ (*PstI* site underlined) and 5’-<u>ggatcc</u>ttctggaaaattattggaaaat-3’(*BamHI* site underlined). The PCR product was cloned into pCR®2.1-TOPO® (Invitrogen), cut out with *PstI* and *BamHI* and ligated into pBluescriptKS-. The *kin-29*cDNA and *unc-54* 3’UTR was amplified, cloned, cut and ligated as described for the *Pges-1::kin-29cDNA* construct (see above). The plasmid was injected into *ceh-10;kin-29* at 25 ng/μl together with 2 ng/μl *Pmyo-2::mCherry*.

***Phlh-1::GFP*** was generated by cutting out *Phlh-1* from pCR®2.1-TOPO® (Invitrogen) (see above) with *PstI* and *BamHI* and ligate the fragment into pPD95.77. The plasmid was injected into wild-type worms at 25 ng/μl together with 40 ng/μl *pRF4 (rol-6 su1006).*

***Pelt-3:: kin-29cDNA*** was generated by PCR amplifying 1964 bp of the *elt-3* promoter using wild type genomic DNA as template with the following primers: 5’-<u>ctgcag</u>tgtgacacgttgtttcacggtc-3’ (*PstI* site underlined) and 5’-<u>ggatcc</u>gaagtttgaaataccaggtagc-3’(*BamHI* site underlined). The PCR product was cloned into pCR®2.1-TOPO® (Invitrogen), cut out with *PstI* and *BamHI* and ligated into pBluescriptKS-. The *kin-29*cDNA and *unc-54* 3’UTR were amplified, cloned, cut and ligated as described for the *Pges-1::kin-29cDNA* construct (see above). The plasmid was injected into *ceh-10;kin-29* at 25 ng/μl together with 2 ng/μl *Pmyo-2::mCherry*.

***Pelt-3::GFP*** was generated by cutting out *Pelt-3* from pCR®2.1-TOPO® (Invitrogen) (see above) with *PstI* and *BamHI* and ligate the fragment into pPD95.77. The plasmid was injected into wild-type worms at 25 ng/μl together with 40 ng/μl *pRF4 (rol-6 su1006)*.

***Prab-3:: kin-29cDNA*** was generated by PCR amplifying 1329 bp of the *rab-3* promoter using wild type genomic DNA as template with the following primers: 5’-<u>ctgcag</u>cgaagctataatagtttttc-3’ (*PstI* site underlined) and 5’-<u>ggatcc</u>ggtcttcttcgtttccgcc-3’(*BamHI* site underlined). The PCR product was cloned into pCR®2.1-TOPO® (Invitrogen), cut out with *PstI* and *BamHI* and ligated into pBluescriptKS-. The *kin-29*cDNA and *unc-54* 3’UTR was amplified, cloned, cut and ligated as described for the *Pges-1::kin-29cDNA* construct (see above). The plasmid was injected into *ceh-10;kin-29* at 10 ng/μl together with 2 ng/μl *Pmyo-2::mCherry*.

***Prab-3::GFP*** was generated by cutting out *Prab-3* from pCR®2.1-TOPO® (Invitrogen) (see above) with *PstI* and *BamHI* and ligate the fragment into pPD95.77. The plasmid was injected into wild-type animals at 10 ng/μl together with 40 ng/μl *pRF4 (rol-6 su1006)*.

***Pkin-29::kin-29cDNA*** was generated by PCR amplifying 1400 bp of the *kin-29* promoter using wild type genomic DNA as template with the following primers: 5’-<u>ctgcagctattactgtaacacctcttac</u>-3’ (*PstI* site underlined) and 5’-<u>ggatcc</u>tgcagtgttggtgtggcggcgc-3’(*BamHI* site underlined). The PCR product was cloned into pCR®2.1-TOPO® (Invitrogen), cut out with *PstI* and *BamHI* and ligated into pBluescriptKS-. The *kin-29*cDNA and *unc-54* 3’UTR were amplified, cloned, cut and ligated as described for the *Pges-1::kin-29cDNA* construct (see above). The plasmid was injected into wild type worms at 25 ng/μl together with 2 ng/μl *Pmyo-2::mCherry*.

***Pkin-29::kin-29::GFP*** was generated by cutting out *Pkin-29* from pCR®2.1-TOPO® (Invitrogen) (see above) with *PstI* and *BamHI* and ligate the fragment into pPD95.77. The plasmid was injected into wild-type animals at 25 ng/μl together with 40 ng/μl *pRF4 (rol-6 su1006).*

***Pkin-29::kin-29SER517ALA*** was generated by modifying *Pkin-29::kin-29cDNA* using PCR-based mutagenesis (Quickchange II XL site-directed mutagenesis kit, Stratagene). The following primers were used: 5’-ccaaagagtgaacgccgagctgccgccggtgaaactcttctgcc-3’ and its reverse complement. The plasmid was injected into wild type worms at 25 ng/μl together with 2 ng/μl *Pmyo-2::mCherry*.

***Pceh-23_L::acy-2 fragment (sense/anti-sense)*** *Pceh-23_L::acy-2 fragment(sense)* was generated with a Gibson assembly cloning kit (NEB) by assembly of the following two DNA fragments: (1) *Pceh-23_L* that was amplified from *Pceh-23_L::unc-53cDNA* with the primers: 5’-ggtactccagccgactccatatgattgcggccgcattttcaaattttaaata-3’and 5’-ctactctccctgtttccagcttatggctgcagtttttctaccggtaccctca-3’and (2) 1252 bp *acy-2* genomic fragment that was amplified from N2 genomic DNA with the primers: 5’-tgagggtaccggtagaaaaactgcagccataagctggaaacagggagagtag-3’and 5’-tatttaaaatttgaaaatgcggccgcaatcatatggagtcggctggagtacc-3’.

***Pacy-2::acy-2 genomic*** was generated with a Gibson assembly cloning kit (NEB) by assembly of the following two DNA fragments: (1) 1200 bp *Pacy-2* that was amplified from wild-type genomic DNA template with the primers: 5’-gctgtctactgccaaatacgtc-3’and 5’-tgcgcgcctggaattcagg-3’and (2) *acy-2* genomic backbone that was amplified from *Pceh-23_L::acy-2(genomic)* with the primers: 5’-cctgaattccaggcgcgcaatgtcgacagtgatggaaatgtcgacg-3’and 5’-gacgtatttggcagtagacagccccagcttttgttccctttagtg-3’. *Pacy-2::acy-2 genomic* was injected into wild-type worms at 20 ng/μl with 3 ng/μl *Pmyo-2::GFP*.

*Pceh-23_L::acy-2 fragment(anti-sense)* was generated with a Gibson assembly cloning kit (NEB) by assembly of the following two DNA fragments: (1) *Pceh-23_L* that was amplified from *Pceh-23_L::unc-53cDNA* with the primers: 5’-ctactctccctgtttccagcttatgggcggccgcattttcaaattttaaata-3’and 5’-ggtactccagccgactccatatgattctgcagtttttctaccggtaccctca-3’and (2) 1252 bp *acy-2* genomic fragment that was amplified from N2 genomic DNA with the primers: 5’-tgagggtaccggtagaaaaactgcagaatcatatggagtcggctggagtacc-3’and 5’-tatttaaaatttgaaaatgcggccgcccataagctggaaacagggagagtag-3’.

*Pceh-23::acy-2 fragment(sense)* and *Pceh-23::acy-2 fragment(anti-sense)* were injected together into wild type worms at 20 ng/μl each with 3 ng/μl *Pmyo-2::GFP*.

***Pceh-23_L::acy-2(genomic)*** was generated with a Gibson assembly cloning kit (NEB) by assembly of the following two DNA fragments: (1) *Pceh-23_L* that was amplified from *Pceh-23_L::GFP* with the primers: 5’-gacactccaaaattttccaaacttaacttataaatcaaaagaatagaccgaga-3’and 5’-cgtcgacatttccatcactgtcgacatctgcagtttttctaccggtaccctca-3’and (2) *acy-2* genomic DNA that was amplified from N2 genomic DNA with the primers: 5’-tgagggtaccggtagaaaaactgcagatgtcgacagtgatggaaatgtcgacg-3’and 5’-tctcggtctattcttttgatttataagttaagtttggaaaattttggagtgtc-3’. *Pceh-23_L::acy-2(genomic)* was injected into wild type worms at 20 ng/μl together with 3 ng/μl *Pmyo-2::GFP*.

Germline transformation was performed by direct injection of various plasmid DNAs into the gonads of adult wild-type animals as described (MELLO *et al.* 1991).

### Survival assay

Eggs were transferred to fresh NGM plates and allowed to hatch. The newly hatched L1 larvae were transferred to new plates and the stage of the worms were studied after 24, 48 and 72 hrs. At least 3 biological replicates have been performed for each strain. Error bars show the 95% confidence interval determined by Z-tests. The p-values are calculated using Fisher’s exact test.

### cAMP feeding

8-Br-cAMP (Tocris) was mixed with fresh growing *E.coli* (OP50) bacteria (grown over night in Luria Broth media with shaking at 37°C). 75 μl of bacteria-cAMP mix was seeded onto small NGM plates and the plates were allowed to dry for 1 h. Worms were transferred to the plates and immediately another batch of 75 μl bacteria-cAMP mix was added on top of the worms. For survival studies, eggs from mothers previously grown on cAMP plates were transferred to new cAMP plates and were allowed to hatch and develop for 72-96 hrs and then the stage of the worms were determined.

### Fluorescence microscopy

Worms were anesthetized in 10 mM levamisole. A Zeiss Axioskop 2 microscope was used to examine worms. Images were collected using an ORCA-ER CCD camera (Hamamatsu) and Openlab imaging software (Improvision).

## Results

### Mutations in kin-29 rescued phenotypes caused by defective CANs

The two bilaterally symmetric *C. elegans* CANs are generated in the head and migrate toward the tail to occupy positions near the center of the embryo (SULSTON 1983). Each CAN extends an anterior axon to the head and a posterior axon to the tail (WHITE *et al.* 1986; WU *et al.* 2011) (Figure 1A). Normal morphogenesis and larval development require CAN neuron function (FORRESTER AND GARRIGA 1997; FORRESTER *et al.* 1998). In mutants lacking *vab-8* function, for example, the CAN cell bodies usually fail to migrate and either lack or have short posterior axons (HEDGECOCK *et al.* 1987; MANSER AND WOOD 1990; WIGHTMAN *et al.* 1996; ADLER *et al.* 2006) (Figure 1C). The lack of CAN function in the posterior of *vab-8* mutants is thought to result in thinning of the posterior body, the Withered tail (Wit) phenotype (Figure 1D). In *ceh-10(gm58)* mutants, the CANs cannot be detected using Nomarski optics or a CAN differentiation marker, and the worms arrest as early larvae (Fig 1*G-H*) (FORRESTER AND GARRIGA 1997; FORRESTER *et al.* 1998). Because laser killing of the CANs also results in a larval arrest (FORRESTER AND GARRIGA 1997), the developmental arrest phenotype of *ceh-10* mutants is thought to result from loss of CAN function. It is unclear, however, whether the CANs are absent in *ceh-10(gm58)* mutants or whether they are present but fail to differentiate.

**Figure 1.**
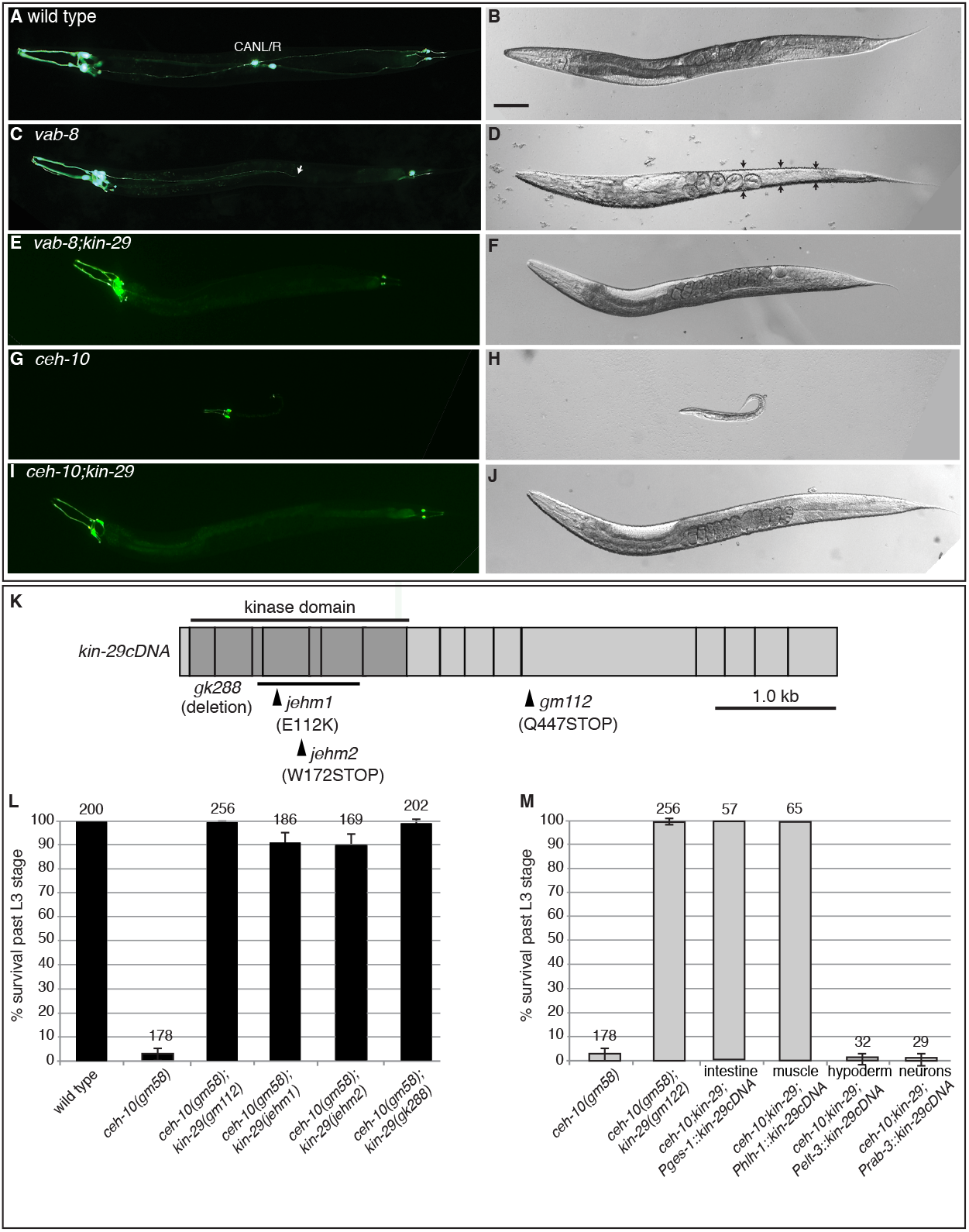
Mutations in *kin-29* rescue phenotypes caused by defective CANs. (A, C, E, G) Fluorescence photomicrographs and (B, D, F, I) Nomarski microscopy of worms containing the *Pceh-23::GFP* transgene, which is expressed in the CANs and in tail and head neurons. (A-J) Anterior is to the left and dorsal is up, the scale bar represents 100 μm. (A) In wild-type worms, the CAN cell bodies are located in the middle of the worm and each neuron extends axons both anteriorly and posteriorly. (B) The body morphology of a wild-type worm. (C) In *vab-8* mutants, the CANs fail to migrate and are located in the head among the other neurons that express GFP. The CAN axons fail to extend to the tail (the arrowhead indicates where one of the axon’s projection stops). (D) The posterior body, as indicated by the arrows, is much thinner in *vab-8* mutants (the Withered Tail or Wit phenotype). (E) In *vab-8; kin-29* double mutants, the CAN migration and extension defects are not rescued, (F) but the Wit phenotype is rescued. (G) In *ceh-10* mutants, the CANs fail to express *Pceh-23::GFP*. (H) *ceh-10* mutants arrest their development during the L1-L2 larval stage. (I) In *ceh-10; kin-29* double mutants, the CANs still are undetectable, (J) but the larval arrest is rescued. (K) The structure of *kin-29* cDNA and the different mutant alleles used in this study. (L and M) Quantification of survival past the L3 larval stage. The number of animals scored for each genotype is indicated above each bar. Error bars show the 95% confidence interval determined by Z-tests. (L) *kin-29* mutant alleles rescue the larval arrest phenotype of *ceh-10* mutants. (M) Tissue-specific expression of *kin-29* cDNA in *ceh-10; kin-29* mutants. *kin-29* cDNA was expressed from an intestinal specific (*Pges-1*), a body wall muscle specific (*Phlh-1*), a hypodermal specific (*Pelt-3*) and a pan-neuronal promoter (*Prab-3*). The number of animals for *ceh-10; kin-29; Pelt-3::kin-29* and *ceh-10; kin-29; Prab-3::kin-29* were small because these animals arrested development and could not be propagated. The arrested transgenic animals were identified based on the presence of the co-transforming marker.

To investigate how the CANs regulate larval development, we carried out two suppressor screens. In the first screen, we mutagenized *vab-8(e1017)* mutants and screened for suppressor mutations that rescued the Wit phenotype without rescuing the CAN migration or axon extension defects and identified *kin-29(gm112).* Because this mutation also suppressed the larval arrest phenotype of *ceh-10* mutants (see below), we mutagenized *ceh-10(gm58)* mutants and screened for mutations that suppressed the larval arrest phenotype but did not restore the CANs based on our inability to detect the cells using Nomarski optics or a *Pceh-23::gfp* reporter transgene. In this screen, we isolated four suppressed strains. Two of these strains contained the *kin-29(jehm1)* or *kin-29(jehm2)* mutations (Figure 1, E-F, I-J).

*kin-29* encodes a serine/threonine kinase that is homologous to the Salt-Inducible Kinases (SIKs) that are related to the AMPK/SNF1 family of kinases (LANJUIN AND SENGUPTA 2002). Sequencing of the *kin-29* gene from the different mutants revealed that the *jehm1* allele is a missense mutation that changes a conserved glutamate in the kinase domain to lysine (E112K), that the *jehm2* allele is a nonsense mutation that changes a conserved tryptophan in the kinase domain to an amber stop codon (W172STOP), and that the *gm112* allele is a nonsense mutation that changes a glutamine to an amber stop codon (Q447STOP) (Figure 1K). We also analyzed the *kin-29(gk288)* allele isolated by the International *C. elegans* Gene Knockout Consortium. The 575 bp deletion removes most of the kinase domain and results in a downstream frameshift. All of the *kin-29* mutant alleles rescued the *vab-8* Wit phenotype (data not shown), and by scoring the ability of animals to develop past the third larval (L3) stage, all of the *kin-29* mutant alleles also suppressed the *ceh-10* mutant larval arrest phenotype (Figure 1L). These findings indicate that the morphological and larval arrest phenotypes caused by CAN dysfunction or loss require *kin-29* function. In all of the studies described below, we used the *kin-29(gm112)* allele.

### kin-29 functions in neurons and hypodermal cells to mediate CAN function

To determine where the *kin-29* mutations act to suppress the *ceh-10* larval arrest phenotype, we expressed a *kin-29* cDNA from cell-specific promoters in *ceh-10; kin-29* double mutants and asked whether *kin-29* expression in specific cell types produced the larval arrest phenotype of the *ceh-10* single mutant. We tested expression in intestine, body-wall muscle, hypodermal cells and neurons, cell types known to express *kin-29* (MADUZIA *et al.* 2005). For intestinal expression we used the *ges-1* promoter (AAMODT *et al.* 1991), for body-wall muscle expression we used the *hlh-1* promoter (QADOTA *et al.* 2007), and for hypodermal expression we used the *elt-3* promoter (GILLEARD *et al.* 1999). For neuronal expression we used the *rab-3* promoter, which is expressed in all neurons except the CANs (STEFANAKIS *et al.* 2015). To confirm that the promoters used to drive *kin-29* in these cells were indeed specific, we also fused the promoters to the GFP gene, studied the expression of the transgenic animals at different developmental stages and found that the promoters drove expression in the predicted cells (data not shown). Only when neurons or hypodermal cells expressed the *kin-29* cDNA was the *ceh-10* larval arrest phenotype restored, suggesting that deregulated *kin-29* activity in either neurons (other than the CANs) or hypodermal cells is sufficient to arrest larval development (Figure 1M).

Because all known CAN promoters require *ceh-10* function, we were unable to confirm that *kin-29* does not act in the CANs to suppress *ceh-10* lethality. However, we were able to express a *kin-29* cDNA in the CANS of *vab-8; kin-29* double mutants since the CANs are present in these animals. To ensure specific expression in the CANs, we used a part of the *ceh-23* promoter that drives expression only in the CANs (*Pceh-23_L*) (WENICK AND HOBERT 2004). CAN-specific expression of *kin-29* did not restore the Wit phenotype (N=50), consistent with the hypothesis that KIN-29 acts neurons and the hypodermis to mediate the effects of CAN function.

### Loss of the MEF-2 MADS domain transcription factor also rescues ceh-10 mutant larval arrest

In *C. elegans* and cultured cells, SIKs phosphorylate and inhibit class II histone deacetylases, which act either upstream of or in a complex with the transcription factor MEF2 to regulate gene transcription (MISKA *et al.* 1999; LANJUIN AND SENGUPTA 2002; CHAN *et al.* 2003; VAN DER LINDEN *et al.* 2008; COHEN *et al.* 2009). Loss of the *C. elegans* homologs *of MEF2* (*mef-2)* and the class IIa HDACs (*hda-4)* suppress several *kin-29* mutant phenotypes: small body size, long lifespan, slow growth, hyper-foraging and chemoreceptor gene regulation (VAN DER LINDEN *et al.* 2007). If the sole activity of KIN-29 in suppressing the larval arrest phenotype of *ceh-10* mutants is to inhibit the function of a HDA-4/MEF-2 repressive complex, then *hda-4* and *mef-2* mutants should exhibit a larval arrest phenotype similar to *ceh-10* mutants, but both *hda-4* and *mef-2* mutants are viable and fertile.

To determine if HDA-4 and MEF-2 function differently in the regulation of morphogenesis and larval development, we asked whether mutations in these genes interacted with a *ceh-10* mutation. Although an *hda-4* mutation had no effect on the *ceh-10* larval arrest phenotype, both the *mef-2(gv1)* mutation and *mef-2(RNAi)* suppressed larval arrest (Figures 1L and 2). The *ceh-10; mef-2; kin-29* triple mutant has survival rates similar to the *ceh-10; kin-29* and *ceh-10; mef-2* double mutants. These findings indicate that the functional relationship between *kin-29* and *mef-2* differs in chemoreceptor regulation and CAN-dependent larval development.

**Figure 2.**
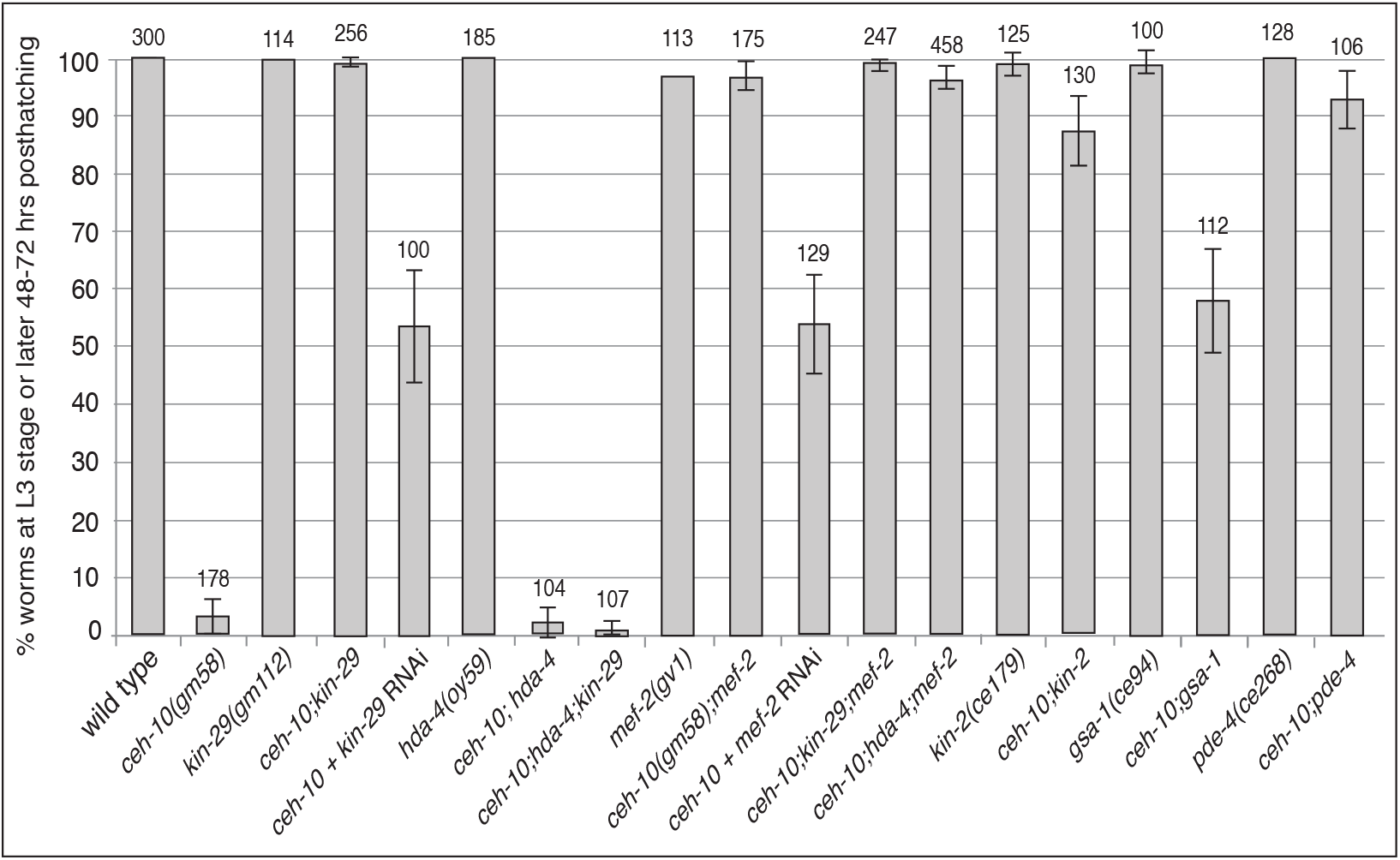
Mutations that upregulate cAMP levels and PKA activity and reduce the function transcription factor MEF-2 rescue the *ceh-10* mutant larval arrest phenotype. Quantification of survival past the L3 larval stage of wild type, single, double and triple mutant strains containing the *ceh-10* mutation. The number of animals scored for each genotype is indicated above each bar. Error bars show the 95% confidence interval determined by Z-tests.

### Mutations that upregulate the cAMP-dependent PKA pathway suppressed ceh-10 mutant larval arrest

cAMP-dependent protein kinase A (PKA) inhibits KIN-29 and its SIK homologs at both the transcriptional and post-translational levels (TAKEMORI *et al.* 2002; OKAMOTO *et al.* 2004; BERDEAUX *et al.* 2007; VAN DER LINDEN *et al.* 2008; WANG *et al.* 2008). In *C. elegans*, PKA consists of two subunits, the catalytic subunit KIN-1 and the regulatory subunit KIN-2. PKA is activated by cAMP that is produced by adenylyl cyclases. One of these, ACY-1, can be activated by the heterotrimeric G protein GSA-1 (BERGER *et al.* 1998). Loss of *acy-1*, *gsa-1*, *kin-1*, and *kin-2* result in embryonic or larval lethality. If the proteins encoded by these genes inhibit KIN-29 function, then like loss-of-function mutations in *kin-29,* gain-of-function mutations in these genes might also suppress the larval arrest phenotype of *ceh-10* mutants. *kin-2(ce179)* mutants express a PKA holoenzyme that is hypersensitive to low levels of cAMP, and *gsa-1(ce94)* mutants express a Gα protein that constitutively activates PKA (KORSWAGEN *et al.* 1997; SCHADE *et al.* 2005; CHARLIE *et al.* 2006). Similar to the *kin-29* loss-of-function mutations, the gain-of-function *kin-2(ce179)* and *gsa-1(ce94)* mutations suppressed the *ceh-10* larval arrest phenotype without suppressing the CAN defects of *ceh-10* mutants (Figure 2 and data not shown).

We also asked whether elevating the levels of cAMP could suppress the *ceh-10* larval arrest phenotype. The gene *pde-4* encodes a cAMP phosphodiesterase that is homologous to human cAMP phosphodiesterase 4D (CHARLIE *et al.* 2006). cAMP phosphodiesterases convert cAMP to 5’-AMP and thus lower cAMP levels (SUNAHARA *et al.* 1996). The *pde-4(ce268)* mutation disrupts the PDE-4 catalytic domain and reduces PDE-4 function, which is predicted to increase cAMP levels (CHARLIE *et al.* 2006). This mutation suppressed *ceh-10* mutant lethality (Figure 2).

### Hyperactive KIN-29 results in larval arrest

We asked if we could phenocopy the larval arrest phenotype of *ceh-10* mutants by introducing a hyperactive version of KIN-29 into wild-type hermaphrodites. We created a construct with a mutation in the conserved PKA phosphorylation site (Ser 517-Ala) (TAKEMORI *et al.* 2002; VAN DER LINDEN *et al.* 2008) and expressed it using the *kin-29* promoter, *Pkin-29::kin-29cDNA(Ser517-Ala*). As controls, we made two constructs lacking the mutation, *Pkin-29::kin-29cDNA* and *Pkin-29::GFP.* The control constructs were injected into wild-type worms, creating several stable transgenic lines lacking obvious phenotypes. The construct with the *kin-29* promoter driving GFP showed the same expression pattern as described previously (LANJUIN AND SENGUPTA 2002). We injected the mutated construct into wild-type hermaphrodites, and 72 % (N=58) of the transgenic worms arrested as early larvae. The surviving transgenic worms failed to produce lines. This finding is consistent with the hypothesis that KIN-29 is a PKA target.

### ACY-2 functions in the CANs to produce essential levels of cAMP

Our results suggest that the CANs signal to the hypodermis and other neurons, activating PKA in these tissues. PKA represses KIN-29, which allows larval development to proceed. Because the *pde-4* mutation rescued the *ceh-10* larval arrest phenotype to a similar degree as the *kin-29* mutations, we asked if exogenous cAMP could also rescue *ceh-10* loss. To explore this possibility, we fed *ceh-10* mutants a synthetic version of cAMP, 8-Br-cAMP, which is a cell-permeable cAMP analog that is resistant to hydrolysis by phosphodiesterases (SANDBERG *et al.* 1991). We tested different concentrations and found that 5 mM 8-Br-cAMP gave the best rescue with 75% of the *ceh-10* mutants developing past the L3 stage (Figure 3). The *ceh-10* mutants could be maintained for generations on 8-Br-cAMP but arrested development if they were moved back to media lacking it (data not shown). cAMP is synthesized from ATP by adenylyl cyclases (ACYs). *C. elegans* has four ACYs: ACY-1, ACY-2, ACY-3 and ACY-4. With the exception of ACY-1, which is broadly expressed in neurons and body wall muscles (MOORMAN AND PLASTERK 2002), the other three ACYs have relatively restricted expression patterns. For example, ACY-2 is only expressed in a few neurons in head ganglia and in the CANs (KORSWAGEN *et al.* 1998) (Table 1). Loss-of-function mutations in *acy-1* or *acy-2* result in larval arrest (KORSWAGEN *et al.* 1998; MOORMAN AND PLASTERK 2002), which prompted us to examine if *acy-1* and *acy-2* mutants arrest development in a similar way to *ceh-10* mutants. The *acy-1(pk1279), acy-2(pk465)* and *ceh-10(gm58)* mutants were maintained as balanced strains. To score the arrested larvae, we picked newly hatched worms that lacked the balancer chromosome and scored their phenotypes after 72 hrs. We noted that the arrested larvae displayed three different phenotypes: normal (Figure 4A), morphological defective (Figure 4B) or clear (Figure 4C). Most of the *acy-1* arrested worms appeared normal, while *acy-2* and *ceh-10* mutants displayed the morphological defective and clear phenotypes at similar frequencies (Figure 4D). We then asked if *acy-1* and *acy-2* mutants could be rescued by the same mutations and treatments as *ceh-10*. The *pde-4* mutation was previously shown to partly suppress the *acy-1* mutant (CHARLIE *et al.* 2006), and we found that it also rescued the *acy-2* mutant (Figure 4E). Feeding *acy-2* mutants with 8-Br-cAMP also rescued larval arrest (Figure 4E). These findings are not unexpected for an adenylyl cyclase mutant. The *kin-29* or *mef-2* mutations, however, suppressed the *acy-2* but not the *acy-1* mutant phenotypes, consistent with the hypothesis that cAMP produced by ACY-2 negatively regulates KIN-29 to promote larval development (Figure 4E, and data not shown).

**Figure 3.**
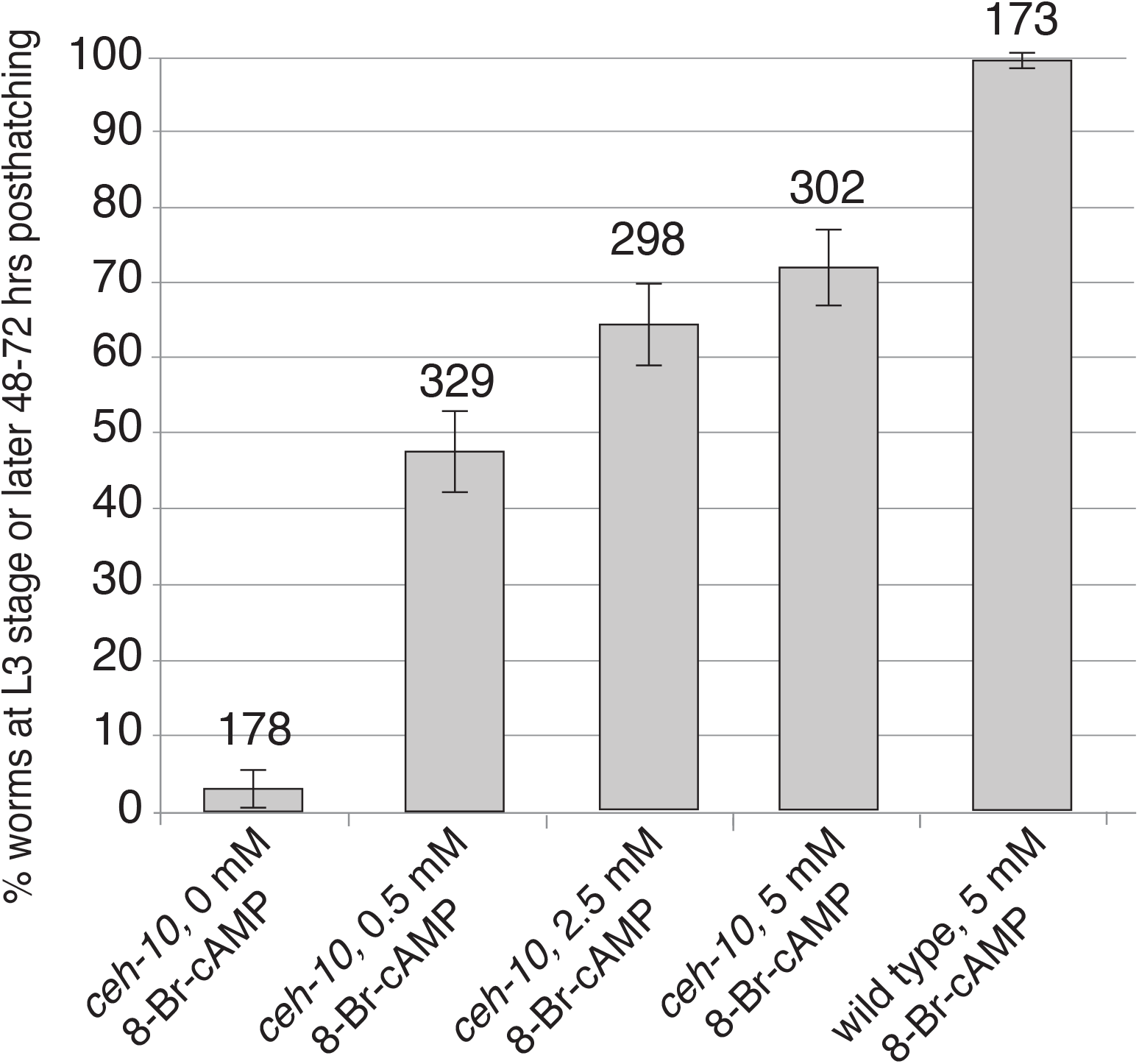
Feeding *ceh-10* mutants synthetic cAMP rescues the larval arrest phenotype. Quantification of survival past the L3 larval stage of *ceh-10* mutants grown on normal plates seeded with bacteria mixed with different concentrations of 8-Br-cAMP. The number of animals scored for each genotype is indicated above each bar. Error bars show the 95% confidence interval determined by Z-tests.

**TABLE 1.**
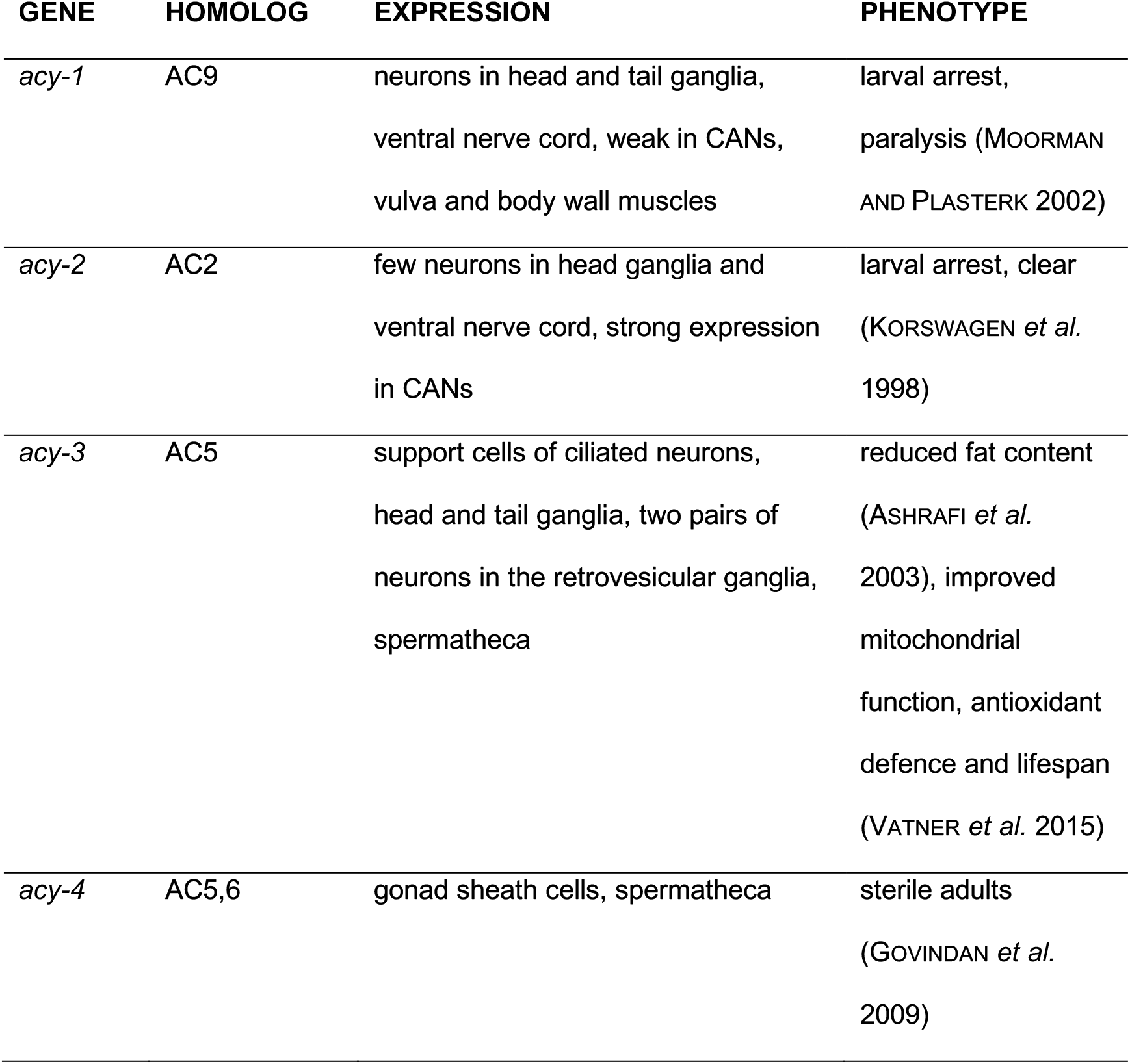
*C. ELEGANS* ADENYLYL CYCLASES.

**Figure 4.**
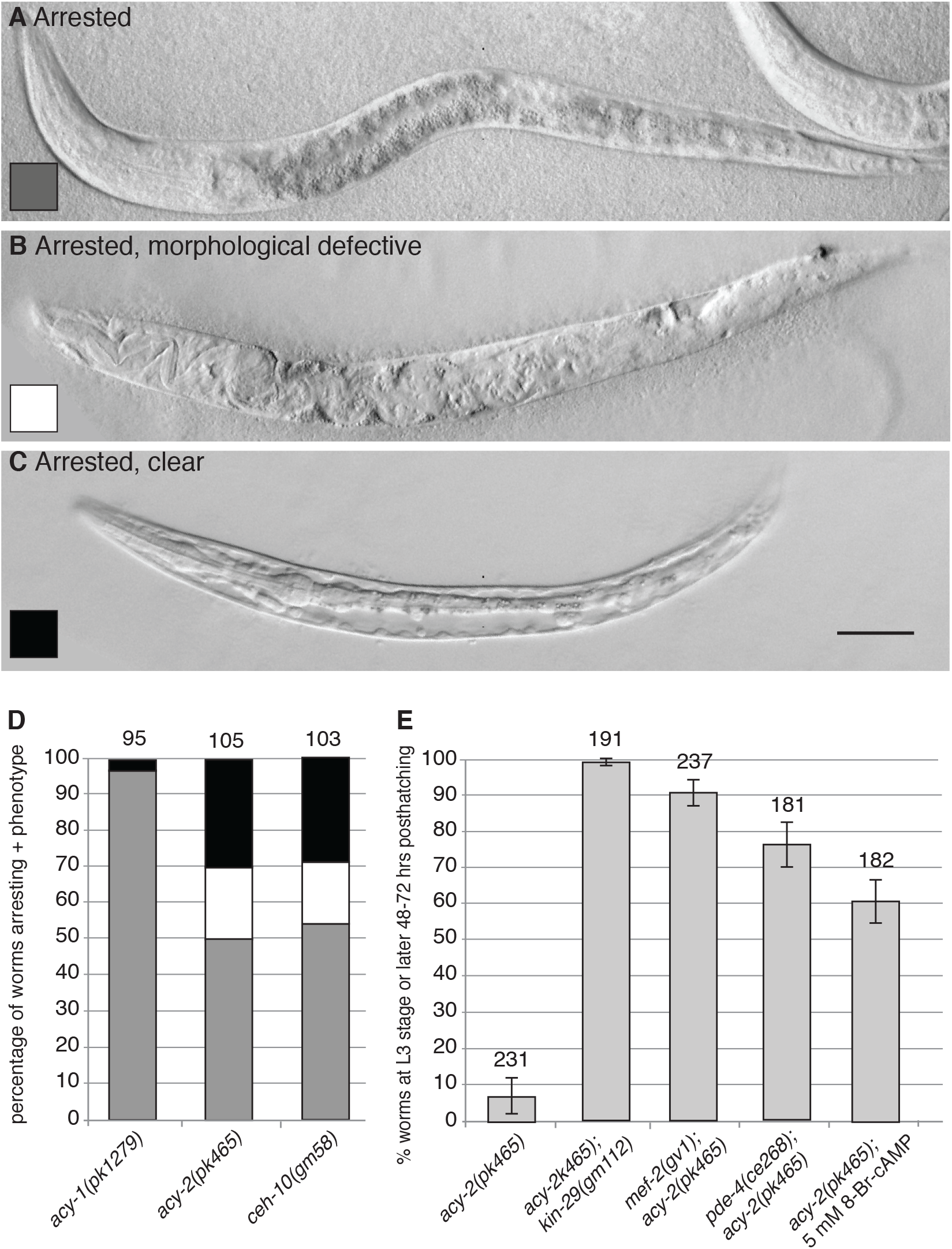
The *kin-29* and *acy-2* mutants have similar phenotypes. (A) An arrested *ceh-10* larva with normal body morphology. The arrowheads mark the width of the intestine. (B) A *ceh-10* mutant with a morphological defective body in which the internal cells appear abnormal. (C) An arrested *ceh-10* mutant with a Clr phenotype. Note that the intestine is much thinner compared to the intestine in the worm in A) (see arrowheads). (D) Quantification of *acy-1*, *acy-2* and *ceh-10* mutants that arrest either with a normal body, a morphological defective body or with a Clr phenotype. (E) Quantification of survival past the L3 larval stage of *acy-2* single mutants, *acy-2; kin-29*, *mef-2; acy-2* and *pde-4; acy-2* double mutants and *acy-2* mutants fed with 5 mM 8-Br-cAMP. The number of animals scored for each genotype in E is indicated above each bar.

Mutations in *kin-29* and *mef-2* rescued the *acy-2* and *ceh-10* mutant defects slightly better than *kin-29* and *mef-2* RNAi (Figures 2, 5A and data not shown). It is noteworthy that many of the RNAi-treated *acy-2* and *ceh-10* mutant worms became visibly sick hours after being transferred to plates with bacteria that did not express *kin-29* or *mef-2* dsRNA (data not shown), suggesting that the activities of KIN-29 and MEF-2 need to be continuously provided for *acy-2* and *ceh-10* worms to survive. These observations imply that CANs need to constantly signal, presumably by producing cAMP that acts in the nervous system and hypodermis.

**Figure 5.**
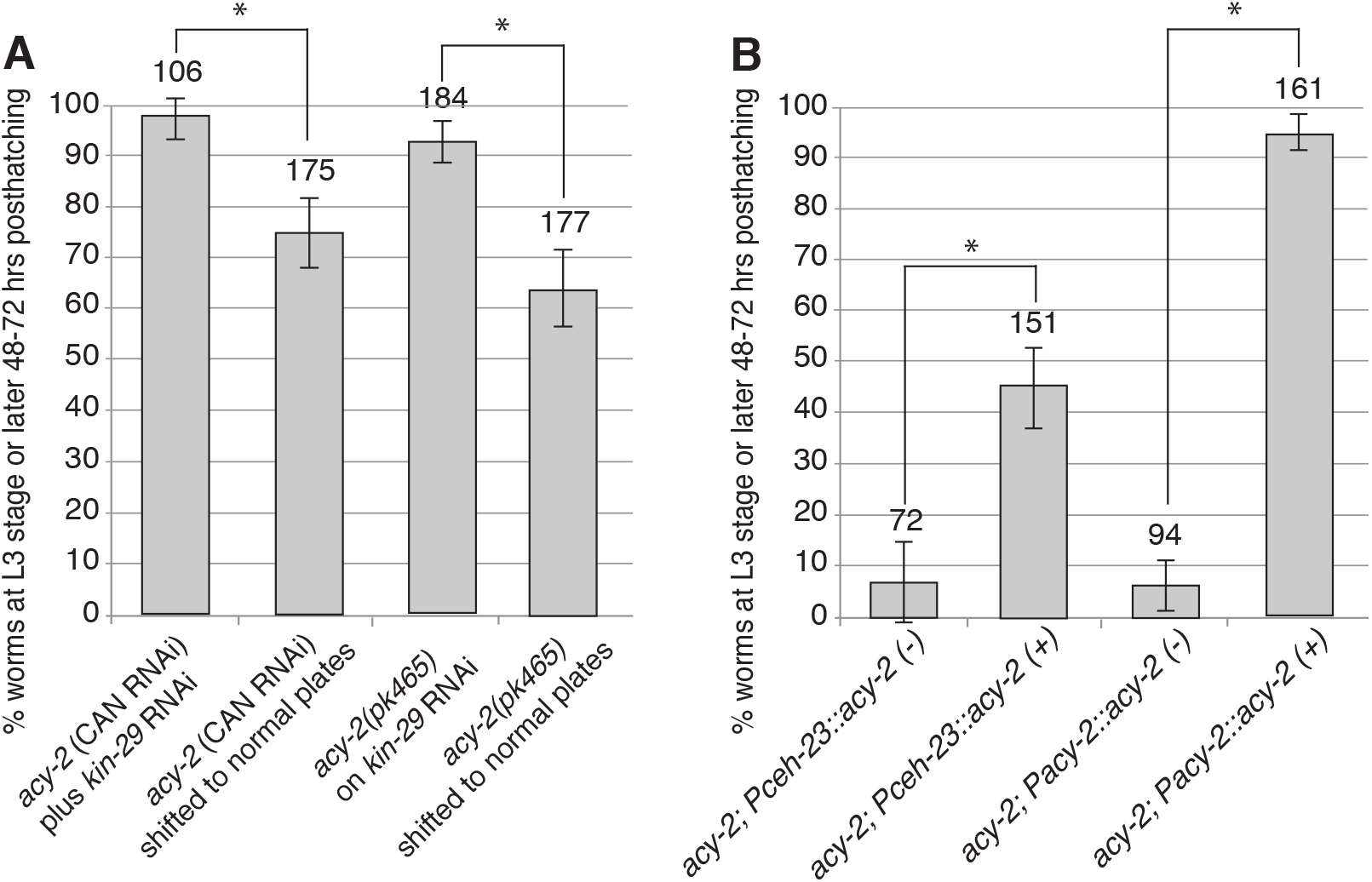
Larval development requires *acy-2* function in the CANs. (A) Quantification of survival past the L3 larval stage of *acy-2* mutant worms or wild-type worms carrying transgenes that express *acy-2* dsRNA specifically in the CANs (*acy-2*(CAN RNAi)). The worms were initially grown on bacteria that produce *kin-29* dsRNA (*kin-29* RNAi) and then transferred to plates with bacteria that do not express *kin-29* dsRNA. (B) Quantification of survival past the L3 larval stage of *acy-2* mutant worms either lacking or carrying the extrachromosomal *PCAN::acy-2* transgene that expresses *acy-2* specifically in the CANs (2 lines). As control ACY-2 was expressed from its endogenous promoter. The number of animals scored for each genotype in E-G is indicated above each bar. Error bars show the 95% confidence interval determined by Z-tests. *: *p*<0.0001 (Fisher’s exact test)

**Figure 6.**
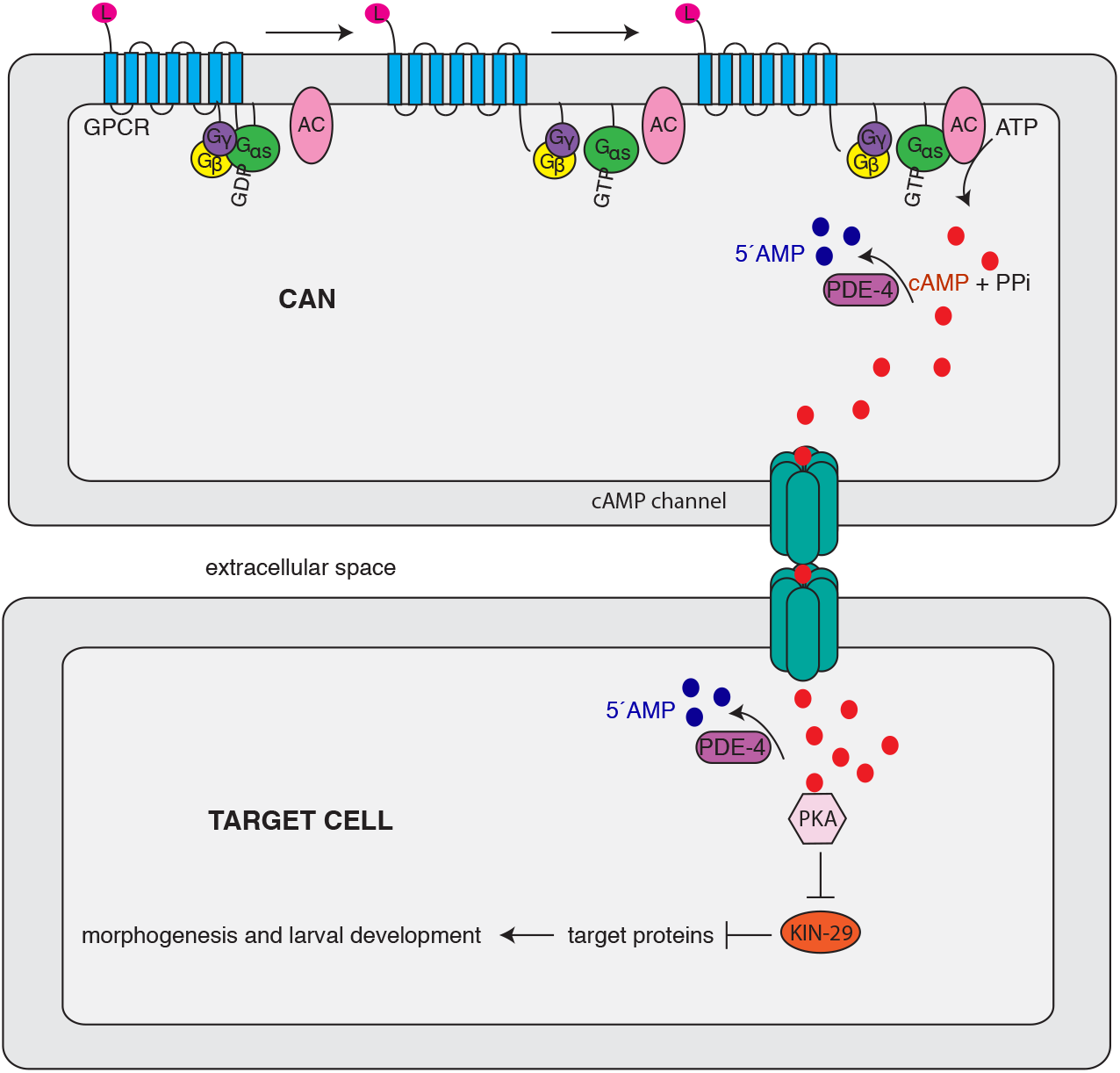
A model for CAN function in larval development. The model proposes that the CANs produce cAMP that diffuses into target hypodermal and neuronal cells where the PKA pathway is activated. A ligand (L) binds to a G protein-coupled receptor (GPCR) located in the cell membrane of the CAN neuron. The GPCR activates the G protein that can in turn activate adenylyl cyclase (AC). AC converts ATP into cAMP that diffuses via cAMP-permeable channels into a target cell. In the target cell cAMP binds to the regulatory subunits of protein kinase A (PKA) that allows dissociation of the catalytic subunits, which then can phosphorylate and inhibit the serine/threonine kinase KIN-29. When KIN-29 is inhibited, KIN-29 target proteins are activated to promote morphogenesis and survival. The levels of cAMP are regulated by phosphodiesterases, for example PDE-4 that degrades cAMP to 5’AMP.

The expression pattern of ACY-2 suggests that it could act in the CANs to promote larval development. To test this hypothesis, we expressed ACY-2 specifically in the CANs of *acy-2* mutants. Our attempt to isolate an *acy-2* cDNA was unsuccessful, possibly because *acy-2* is only expressed in a few cells (KORSWAGEN *et al.* 1998). Instead, we expressed *acy-2* genomic DNA fused to GFP under the control of the CAN-specific promoter *Pceh-23_L* (WENICK AND HOBERT 2004). We generated two independent, non-integrated transgenic lines (*Pceh-23_L::acy-2(genomic)::GFP*), and only the CANs and a single tail neuron expressed these transgenes (not shown). We compared transgenic and non-transgenic worms originating from the same transgenic mother. Both transgenic lines rescued the larval arrest phenotype, with more than 40% of the transgenic worms developing past L3 stage (Figure 5b). We also generated transgenic lines that express *acy-2* from its endogenous promoter (*Pacy-2::acy-2(genomic).* We observed a more robust rescue compared to CAN-specific expression of ACY-2 with more than 90% of the transgenic worms developing past the L3 stage. This finding suggests either that the endogenous promoter drives higher levels of *acy-2* in the CANs or that neurons other than the CANs are also important for *acy-2* mutant larvae to develop.

As an alternative test of this hypothesis, we performed CAN-specific *acy-2* RNAi to ask if we could phenocopy the *acy-2* mutant phenotype. We generated transgenes that expressed both *acy-2* sense and *acy-2* antisense RNA driven from the CAN promoter to generate *acy-2* dsRNA in the CANs. These transgenes were expressed in an *ergo-1* mutant to sensitize the background to RNAi effects (PAVELEC *et al.* 2009). To obtain viable transgenic lines, we grew the worms on plates with bacteria that produced dsRNA that targeted *kin-29,* transferred transgenic worms to plates with control bacteria that did not express *kin-29* dsRNA and scored survival in the next generation. As a control, we subjected *acy-2(pk465)* mutants to the same protocol. When the *Pceh-23_L::acy-2(RNAi)* transgenic worms and the *acy-2* mutants were transferred to plates with control bacteria, both strains arrested development at similar frequencies (Figure 5A). These findings further support the hypothesis that ACY-2 can act in the CANs to promote larval development.

## DISCUSSION

### A conserved pathway mediates the essential function of the CANs

The function of the CANs is mysterious. It has been proposed that the CANs regulate the function of the excretory canal cell, which is involved in osmoregulation (HEDGECOCK *et al.* 1987; FORRESTER AND GARRIGA 1997). This hypothesis is based on the accumulation of fluid in the pseudocoelom, the Clear (Clr) phenotype, in animals missing their CANs. In screens for mutations that bypass the requirement for the CANs in larval development, we identified the gene *kin-29*, which encodes a SIK homolog, and showed that the CANs regulate a conserved cAMP pathway that inhibits *kin-29*.

The KIN-29/SIK pathway mediates diverse functions that range from transcriptional regulation of *C. elegans* chemoreceptors to lipid metabolism in adipocytes (VAN DER LINDEN *et al.* 2007; HENRIKSSON *et al.* 2012). In *C. elegans, kin-29* functions in sensory neurons to regulate body size, entry into the dauer stage, and foraging behavior (LANJUIN AND SENGUPTA 2002; MADUZIA *et al.* 2005). Although the CANs have no obvious role in any of these processes, the essential nature of the cell makes testing its role in other processes difficult. In this context, it is noteworthy that the analogous cell in the nematode *Pristionchus pacificus* expresses the gene *dauerless,* which antagonizes dauer development. Although killing the CAN-like cell in this species does not cause larval arrest, it does cause entry into dauer, presumably due to the lack of *dauerless* expression (MAYER *et al.* 2015).

Suppression of the *ceh-10* larval arrest phenotype by mutations in genes that act in the *kin-29* pathway suggests that the adenylyl cyclase ACY-2 acting through PKA inhibits KIN-29 activity, allowing larval development to progress. It is unclear whether the CANs and this pathway regulate larval development directly or a physiological state that allows development to proceed.

SIKs inhibit the function of class IIa histone deacetylases, which can interact with the MEF2 transcription factor to repress the transcription of target genes (DI GIORGIO AND BRANCOLINI 2016). The sole *C. elegans* member of the class IIa HDAC family is HDA-4. Mutations in either *hda-4* or *mef-2* suppress the effects of *kin-29* mutations on chemoreceptor gene transcription, consistent with the inhibition of HDA-4/MEF-2 repressor activity by KIN-29 (VAN DER LINDEN *et al.* 2007). If KIN-29 inhibits HDA-4/MEF-2 repressor functions in larval development as it does in chemoreceptor regulation, *hda-4* and *mef-2* mutations should cause larval arrest, which they do not. The *mef-2* mutation, however, suppressed the larval arrest phenotype of the *ceh-10* mutant. One model to explain these observations is that KIN-29 activates MEF-2, either directly or indirectly. Stimulation of cortical neurons by BDNF results in the activation of MEF2 transcriptional targets. In these cells, SIK1 phorphorylates the class IIa histone deacetylase HDAC5, resulting in HDAC5 export from the nucleus and allowing MEF2 to function as a transcriptionally activator (FINSTERWALD *et al.* 2013). KIN-29 could also indirectly activate MEF-2 function in larval development, but our genetic results indicate that a protein other than HDA-4 would link KIN-29 to MEF-2 function.

### cAMP transport to target tissues

cAMP has traditionally been described as an intracellular “secondary messenger” that is released in response to signals from “first messengers.” If the hypothesis suggested by our results is correct, how then can cAMP produced in the CANs regulate KIN-29 in other cell types? One interesting possibility is cAMP could diffuse between the CAN and other cells via gap junctions, intercellular channels that allow passive transport of ions and small molecules. Vertebrate gap junctions are hemichannels that consist of connexin (Cx) proteins (ELFGANG *et al.* 1995). Several studies have shown that cAMP diffuses between cells via connexins; for example, it is well established that cAMP passes through the Cx26, Cx32, Cx36, Cx43, Cx45 and Cx47 gap junction channels in Hela cells (BEDNER *et al.* 2003; BEDNER *et al.* 2006; HERNANDEZ *et al.* 2007; CHANDRASEKHAR *et al.* 2013).

In support of this hypothesis, larval development requires gap junction function. *C. elegans* gap junctions are assembled from innexin (INX) proteins (PHELAN *et al.* 1998). The CANs express the innexin genes *inx-3, inx-12* and *inx-13* (ALTUN *et al.* 2009), and RNAi of *inx-3* and mutations in *inx-12* and *inx-13* result in larval arrest (JOHNSEN *et al.* 2000; STARICH *et al.* 2001; SIMMER *et al.* 2003; HALL 2016). CAN-specific RNAi of *inx-13* also produced larval arrest (not shown), which suggests that the gap junctions between the CANs and other cells promote larval development. However, loss of neither *kin-29* nor *mef-2* suppressed the larval arrest phenotype of *inx-12* and *inx-13* mutants (not shown). The lack of suppression may reflect the expression *of inx-12* and *inx-13* in other cell types such as pharynx and muscle, where they could function to promote larval development. Further experiments will be required to test this interesting hypothesis.

## Acknowledgments

The authors thank Dawna Sweeney and Ranjan Devkota for assistance in the lab and valuable discussions. We thank Piali Sengupta for the plasmid containing *Pkin-29::kin-29cDNA::GFP.* Some of the nematode strains used in this study were provided by the *Caenorhabditis Genetics Center*, which is funded by the NIH National Center for Research Resources (NCRR). The *kin-29(gk288)* strain was provided by the *C. elegans* Gene Knockout Facility at the Oklahoma Medical Research Foundation (funded by the National Institutes of Health) and the *C. elegans* Reverse Genetics Core Facility at the University of British Columbia (funded by the Canadian Institute for Health Research, Genome Canada, Genome BC, the Michael Smith Foundation, and the National Institutes of Health). This work was supported by National Institutes of Health grant NS32057 to G.G.

